# Deep Learning for High-Throughput Quantification of Oligodendrocyte Ensheathment at Single-Cell Resolution

**DOI:** 10.1101/389932

**Authors:** Yu Kang T Xu, Daryan Chitsaz, Robert A Brown, Qiao Ling Cui, Matthew A Dabarno, Jack P Antel, Timothy E Kennedy

## Abstract

High-throughput quantification of oligodendrocyte (OL) myelination is a significant challenge that, if addressed, would facilitate the development of therapeutics to promote myelin protection and repair. Here, we established a quantitative high-throughput method to asses OL ensheathment in-vitro, combining nanofiber culture devices and automated imaging with a heuristic approach that informed the development of a deep learning analytic algorithm. The heuristic approach was developed by modeling general characteristics of OL ensheathments, while the deep learning neural network employed a UNet architecture with enhanced capacity to associate ensheathed segments with individual OLs. Reliably extracting multiple morphological parameters from individual cells, without heuristic approximations, mimics the high-level decision-making capacity of human researchers and improves the validity of the neural network. Experimental validation demonstrated that the deep learning approach matched the accuracy of expert-human measurements of the length and number of myelin segments per cell. The combined use of automated imaging and analysis reduces tedious manual labor while eliminating variability. The capacity of this technology to perform multi-parametric analyses at the level of individual cells permits the detection of nuanced cellular differences to accelerate the discovery of new insight into OL physiology.

## 1 Introduction

In the CNS, oligodendrocyte (OLs) produce myelin by wrapping specialized multi-lamellar membrane sheaths around axons. In pathological conditions, like multiple sclerosis, leukodystrophies, or CNS trauma, myelin is progressively lost due to inadequate repair of damaged sheaths [1]. Enhancing myelin production and stability are critical clinical objectives, and several in vitro systems have been established to study these dynamic process [2, 3, 4]. A particularly reduced system cultures OLs on axon-sized electrospun polymer nanofibers. Immature O4+ OLs initially contact fibers, followed by differentiation and expression of mature OL markers like myelin-basic protein (MBP) [5]. The extent of ensheathment around these axon-like structures can be measured in response to different treatments, ranging from substrate coatings to the application of bioactive small molecules [6, 7, 8, 9]. Regardless of the culture system, however, OL myelination is typically quantified manually, creating a significant bottleneck and introducing human subjectivity.

To resolve this quantitative challenge, we developed automated high-throughput methods that combine micro-engineered nanofiber culture devices, automated microscopy, and analytic algorithms to extract detailed morphological properties from individual OLs (Figure 1A). Quantitative algorithms were developed to assess myelin elaborated on electrospun 2 µm-diameter poly-L-lactic acid fibers [6, 7]. OLs cultured on nanofibers recapitulate biologically significant features of myelin development and the planar parallel fiber arrangement is conducive to automated imaging and segmentation. In this report, we describe optimized protocols for OL nanofiber cultures and automated imaging, along with the development of a heuristic algorithm that ultimately informed the construction of a deep learning neural network to quantify immunocytochemically labelled markers of OL ensheathment.

**Figure 1:**
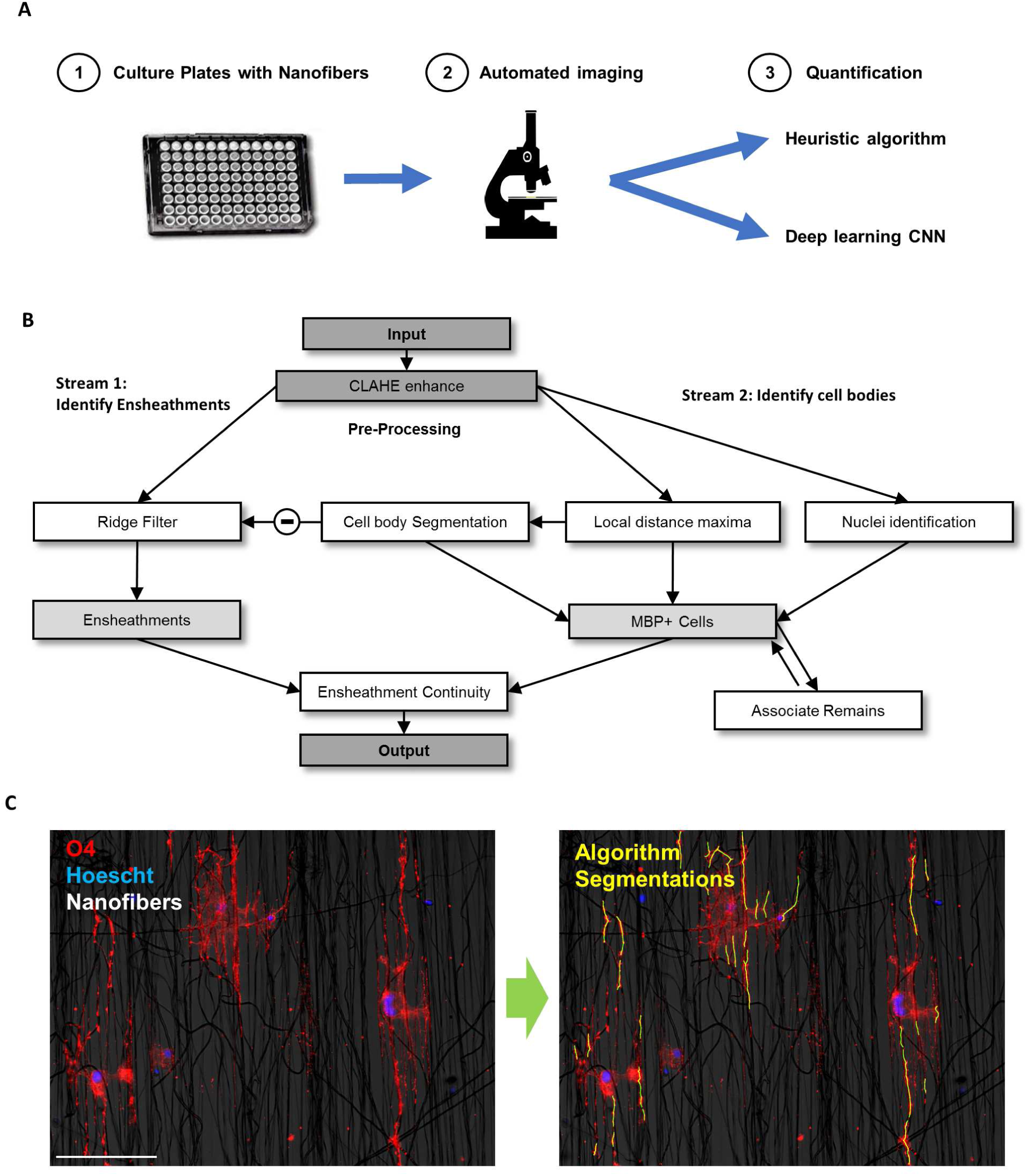
Overview of analytic pipeline and heuristic MATLAB algorithm. (A) Graphic illustrating automated pipeline using nanofiber culture devices, automated imaging, and analytic algorithms to quantify OL ensheathments. (B) Input images are pre-processed using CLAHE enhancement to increase contrast before identifying ensheathments and OL cell bodies in two separate analytic streams. (Stream 1) a ridge-filter is applied to identify ensheathed nanofiber segments and (Stream 2) cell-bodies are estimated using watershed segmentation with imposed local distance maxima. The middle link (subtraction sign) subtracts the cell bodies from the ensheathments identified by the ridge filter to eliminate segments located within cell bodies. The final outputs of both pathways are combined for ensheathment continuity analysis. (C) Input and output of the heuristic algorithmic approach. Cell nuclei (blue), O4 (red), nanofibers (bright field in background). Scale bar 20 *µ*m. Yellow outlines in the output image indicate the segmentations performed by the algorithm.

Our initial “heuristic” approach targeted the general morphological characteristics of OL sheaths, such as the presence of thick elongated structures, to segment fluorescently labelled OL segments. Although the approximations encoded in the MATLAB algorithm resulted in rapid and accurate measurements of individual ensheathed segments, this method revealed that reliably associating myelin-like ensheathments to their corresponding cell bodies presented a substantial challenge. To improve beyond this initial approach, we developed a deep learning neural network with the ability to extract and associate multiple morphological parameters with individual OLs.

To develop the convolutional neural network (CNN), we employed a cell-by-cell training method and a “UNet” architecture [10] that enhanced the algorithm’s capacity to perform segmentations on individual OLs using spatial and object specific information. Validation demonstrated that the deep-learning approach performs beyond the accuracy of the heuristic algorithm, particularly at single-cell resolution. Further, the CNN reached a level of analytic accuracy within the range of human experts on multiple parameters, while reducing workloads and providing standardization that eliminates human variability.

We then tested the sensitivity of the heuristic and CNN algorithms to detect a previously reported difference in the number of sheaths formed per OL on poly-D-lysine (PDL) verses laminin 1 coated nanofibers [7]. Both algorithms successfully detected the difference in number of sheaths and, due to the high number of cells analyzed, also revealed a previously unidentified increase in the length of OL sheaths on laminin 1 compared to PDL coated nanofibers. Our findings highlight the power of this analytic approach to quantify large numbers of cells, thereby providing sufficient sensitivity to detect nuanced differences in cellular processes. Overall, the analytic pipeline we describe allows for a 96 or 384 well plate to be imaged overnight and analyzed the following day with a single computing station. By extracting multiple morphological parameters at single-cell resolution, while obtaining analytic quality similar to human segmentations, this high-throughput system has the potential to accelerate the discovery of therapeutics that promote myelin protection and enhance our understanding of OL physiology.

## 2 Materials and Methods

### 2.1 Electrospun nanofiber OL cell culture and staining

The study was conducted in accordance with the guidelines of the Canadian Council for Animal Care and approved by the McGill University Animal Care Committees. Primary rat OLs were seeded into 96-well plates containing poly-L-lactic acid (PLLA) nanofibers electrospun onto the optically clear base (The Electrospinning Company). Wells were coated with 5 *µ*g/mL of poly-D-lysine (PDL) in PBS for 1 hr at room temperature (RT, approximately 20°C), followed by 2 hr at 37°C with 10 *µ*g/mL recombinant laminin 1 in HBSS (Sigma). OL progenitor cells, isolated via a protocol adapted from McCarthy and de Vellis [11], were seeded at 6000 cells per well in differentiation media, as described in the supplementary methods [12]. Cells were cultured for 7 or 14 days, with full media changes every 2 days or half changes every 4 days, for O4 and MBP-based experiments respectively. After fixation with 4% paraformaldehyde, cells were stained with mouse anti-O4 (RD Systems) or chicken anti-MBP (Aves labs) to mark maturing OLs and their sheaths, as well as Hoescht 33258 and Alexa-546-conjugated phalloidin (Molecular Probes) to mark cell nuclei and processes.

### 2.2 Automated Imaging

Nanofiber plates were imaged using an LSM 880 inverted confocal laser scanning microscope (Carl Zeiss) with a 10X (NA 0.45) Plan-Apochromat objective. Alexa-555-conjugated secondary antibodies (ThermoFisher) were used to ensure minimal spectral overlap so that Hoescht and MBP signals could be acquired on a single track to reduce acquisition time. Acquisitions were automated using Zen Black Systems v2.3 and the Tiles and Positions software module. For each experiment, the user is only required to set the PMT gain to account for differences in staining intensity and to calibrate the motorized stage to target the center of the first well of each plate. In 16 hrs of acquisition, the system imaged ∼4 × 4 frames centered in the middle of each well to cover a ∼4 × 4 mm area with a pixel resolution of 0.5 *µ*m, thereby acquiring >50% of every well with no bias. Focus was performed automatically for each well with the DefiniteFocus.2 module (Carl Zeiss). Since the nanofibers are not perfectly planar, three to five confocal slices covering 10-20 *µ*m in the Z-dimension were acquired for each frame to ensure that all cells were sampled. At lower resolutions, each well took < 10 min to acquire, allowing the whole plate to be imaged in less than 16 hrs. Z-stacks were stitched and compressed via the Extended Depth of Focus (maximum projection) algorithm in Zen Black prior to exporting the images as TIFF files.

### 2.3 Manual segmentation criteria

A set of criteria was defined for classification of OL ensheathments based on morphological characteristics visualized using Hoechst, O4, and MBP staining: (1) The presence of an identifiable cell body and nucleus; (2) The presence of ensheathed processes, defined as segments of O4 or MBP running parallel to the nanofibers in the culture plate having a length > 12 *µ*m and a thickness > 2 *µ*m; (3) Continuity of the cell body with the ensheathments; (4) Each ensheathed fiber may only be associated with 1 cell nucleus. Human researchers were given these criteria and instructed to trace the border of OL ensheathments using the ImageJ polygon tool. These manual segmentations were then used to train and validate the deep learning network.

### 2.4 Heuristic segmentation algorithm

A heuristic algorithm, written in MATLAB, was first constructed to implement the manual segmentation criteria. To fulfill criteria 1, cell bodies were identified through colocalization of Hoechst stained nuclei with MBP-positive cytoplasm. To separate overlapping and crowded cell bodies, the local maxima of a distance transformed MBP image were used for watershed segmentation. For criteria 2, after convolution with second derivative Gaussian filters, ensheathed nanofibers were segmented using a ridge-filter that extracts the eigenvalues from the Hessian matrix of the MBP image. The sensitivity of segmentation was user specified and the diameter of fibers segmented could be modified by altering the sigma of the initial Gaussian filters. The algorithm then cycled through each cell body to associate segmented fibers with corresponding nuclei, thereby satisfying criteria 3 and 4 (Figure 1B). An important limitation of this heuristic algorithm is that the identification of ensheathments and the association of segments to individual cells is performed as two separate quantitative steps. The ridge-filter is applied to extract ensheathments globally from an image and then the local distance maxima of cell bodies is used to associate those segments to individual OLs based on proximity to mimic a cell-by-cell level of quantification (Figure 1B). By separating the analysis into two streams, however, the algorithm does not perform relatively high-level decisions required to reliably associate ensheathments to specific cells when faced with over-crowded images or when fluorescent staining is faded. In contrast, the deep learning approach overcomes these challenges by using spatial and object-specific information simultaneously to analyze and associate OL ensheathments with individual cells.

### 2.5 Neural network training data

To train a neural network to extract morphological features from individual OLs, we generated 3-channel images for our input data, with the middle channel replaced by a mask over a specific nucleus of interest (Figure 2A). This mask prompted the network to learn to associate OL ensheathments with individually masked nuclei. The input data was also appended with a ground-truth image, containing the human tracings of OL ensheathments. To create the ground-truth images, human researchers: (1) manually traced a single cell nucleus with its corresponding ensheathments, saved that as an ROI file in ImageJ, and then repeated the process for every cell in a whole-well image; (2) The ROIs were then used to generate two 16-bit images, one containing all manually traced nuclei, and the other containing all manually traced ensheathments; (3) A Python script then looped through the nuclei-containing image, using each nucleus as a center to crop/create input and ground-truth images for every cell in a full-well image. Cropping input data using individual cell nuclei significantly increased the number of training images, generating ∼40, 000 input examples from a dozen full-well images.

**Figure 2:**
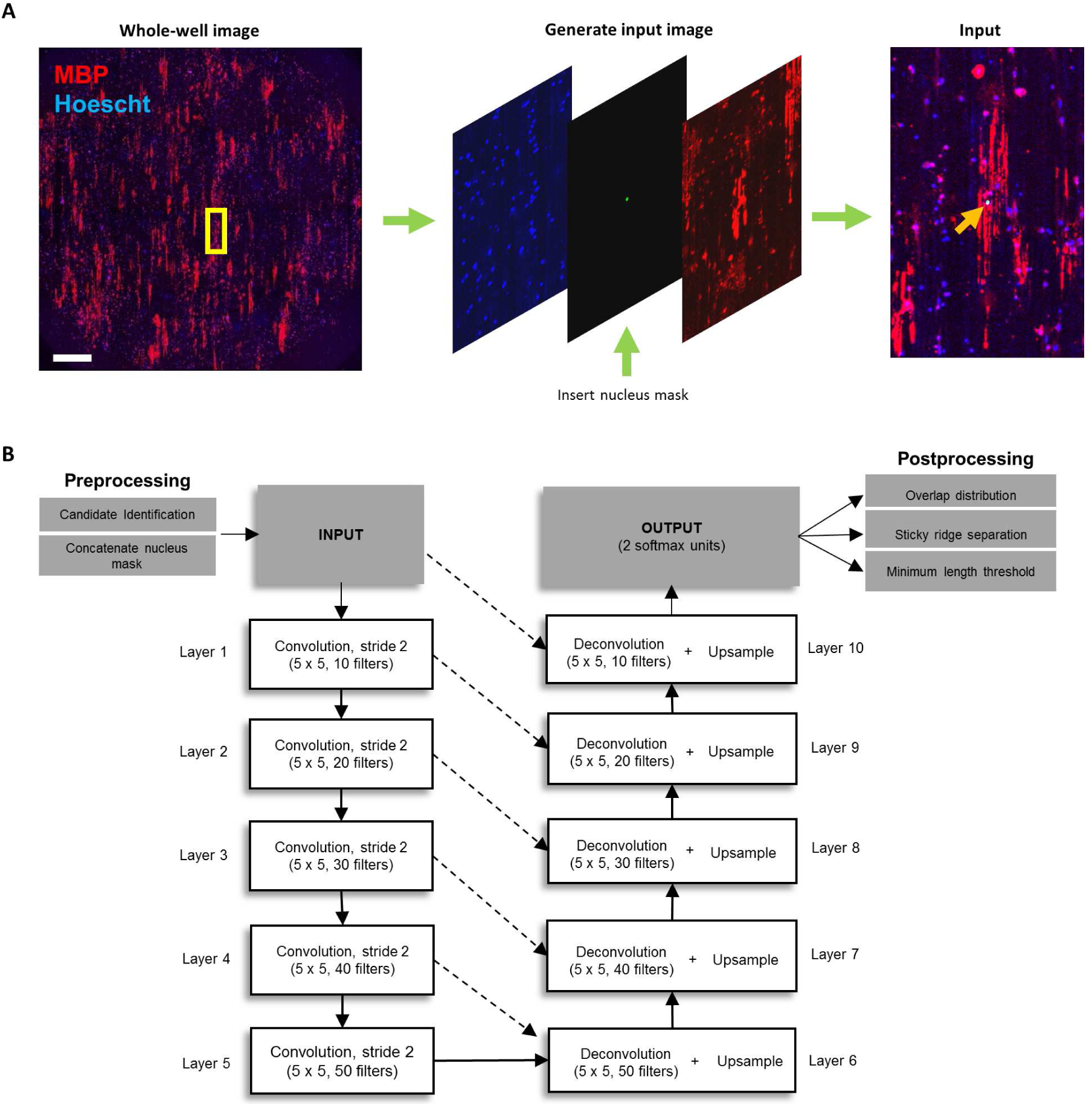
Pre-processing, network architecture and work-flow of CNN approach. (A) The yellow rectangular mask on the whole-well image (left) indicates the portion of the image cropped around a candidate Hoechst center. That Hoechst center is then used as a mask to replace the green channel (middle) in the “Input” image (right). Arrowhead on “Input” image indicates the nucleus of the masked cell of interest. This method is used to generate both the training images and the input images when testing the algorithm. Scale bar corresponds to 1 mm. (B) Starting from the left, a whole-well image is pre-processed to generate nuclei masks for candidate cells. These candidates are then piped into the network architecture as an “input” image that is down-sampled on the left arm of the network, and then up-sampled on the right-arm of the network. Spatial information from the down-sampling branch is also mixed into the up-sampling branch to allow the network to learn to associate ensheathments with individual OLs. The final “output” segmentation is then subjected to further post-processing.

### 2.6 Neural Network Architecture

Google’s open source Tensorflow [13] was used to implement a CNN with a “UNet” architecture [10]. The U-shaped design was selected for its capacity to learn to associate ensheathments with individual OLs, as the down-sampling arm of the network learns features at a variety of scales, and the up-sampling arm combines spatial information from earlier layers to produce a high resolution final segmentation (Figure 2B). The down-sampling branch contained 5 two-dimensional convolutional layers with ∼5 × 5 pixel kernels and a stride of 2. The number of filters at each subsequent layer increased by 10 sequentially from 10 to 50. The up-sampling branch of the network was symmetric to the down-sampling branch but used fractional strides to up-sample the information back to the original size. The up-sampling convolutional layers had ∼5 × 5 pixel kernels and a decreasing number of filters that dropped sequentially by 10 from 50 to 10. The up-sampling branch also mixed in spatial information by concatenating data from earlier layers with similar resolution prior to each convolution. The output of the up-sampling branch was then passed through a softmax convolutional layer with a ∼1 × 1 pixel kernel to generate two mutually exclusive probabilistic output categories corresponding to background and fibers. This probabilistic output was then thresholded at a value of 0.5 to produce a binary segmentation containing the ensheathments identified by the network for a single cell.

### 2.7 Pre-processing data for CNN analysis

An image first undergoes pre-processing by watershed segmentation to identify Hoechst stained nuclei. These nuclei are then used to generate “input” images (Figure 2A). To avoid redundancy, we recognized that some cells are clearly not ensheathed—defined as having zero O4 or MBP near the cell nucleus. Such cells are excluded to save analytic time by binarizing the O4 or MBP channel followed by morphological closing then opening to generate blobs. Only nuclei within O4 or MBP blobs were considered “candidates” for quantification, thereby avoiding analysis of unstained cells.

### 2.8 Post-processing after CNN analysis

After the network cycles through each “candidate” cell within a full-well image, we applied three post-processing steps to assign “overlapped” ensheathments—associated to more than one cell—while removing “sticky” connections (see below) and setting a minimum threshold for ensheathment length (Figure 2B):

1. ***Method of unfair distribution:*** Given the choice to assign an ambiguous “overlapped” ensheathment to a large cell with multiple ensheathed nanofibers, or a small cell with only a single ensheathment, this method operates under the assumption that it is more likely that the uncertain segment will belong to the larger cell.
2. ***Sticky-removal:*** To separate “sticky-segments”—adjacent ensheathments that touch and are wrongly identified as a single object—the method skeletonized the output image and removed all branch points. It then cycled through all the broken segments to eliminate any that were horizontal. Branch points were then re-inserted to restore the vertical connectivity of the ensheathments without their “sticky” horizontal connections.
3. ***User-specified min-length:*** Ensheathments below a minimum length (∼12 *µ*m) were eliminated. As well, since we noticed slight over-sensitivity toward cells with only one or two ensheathments in human non-experts and machines, we also programmed a sensitivity measure to reduce variability across experiments. We recommend a length threshold that is 3–4 times more strict for those single and doubly-ensheathed cells to reduce the chance of identifying false positives, but this value can be adjusted to suit the researcher’s needs. Length was determined by fitting the objects with an ellipse and calculating the distance of the major axis, as defined in regionprops OpenCV [14].

### 2.9 Training

To train a deep learning model to match the cell-by-cell analysis performed by human researchers, we presented the network with ∼40, 000 input images that each contained a masked nucleus of interest (Figure 2A–B). 1,111 of these cropped images were utilized for validation during training, allowing the performance of the model to be followed as it learned. The performance of our CNN in task-learning was monitored through the loss value and Jaccard index (JI). Loss functions serve as a numerical representation of the magnitude of error for a prediction made by the CNN. Here, we used the categorical cross-entropy, on a per-pixel basis, as the loss function to measure the difference between the output of the network and the manually traced ensheathments generated by cell biologists (ground truth). The JI was used to measure the overlap similarity between the binarized output of the CNN and the ground-truth [15]. In our model, a decrease in loss and increase in JI, calculated on the set of 1,111 validation images, served as an indicator of successful task-learning and the generalizability of the network to new data (Figure 3A-B).

**Figure 3:**
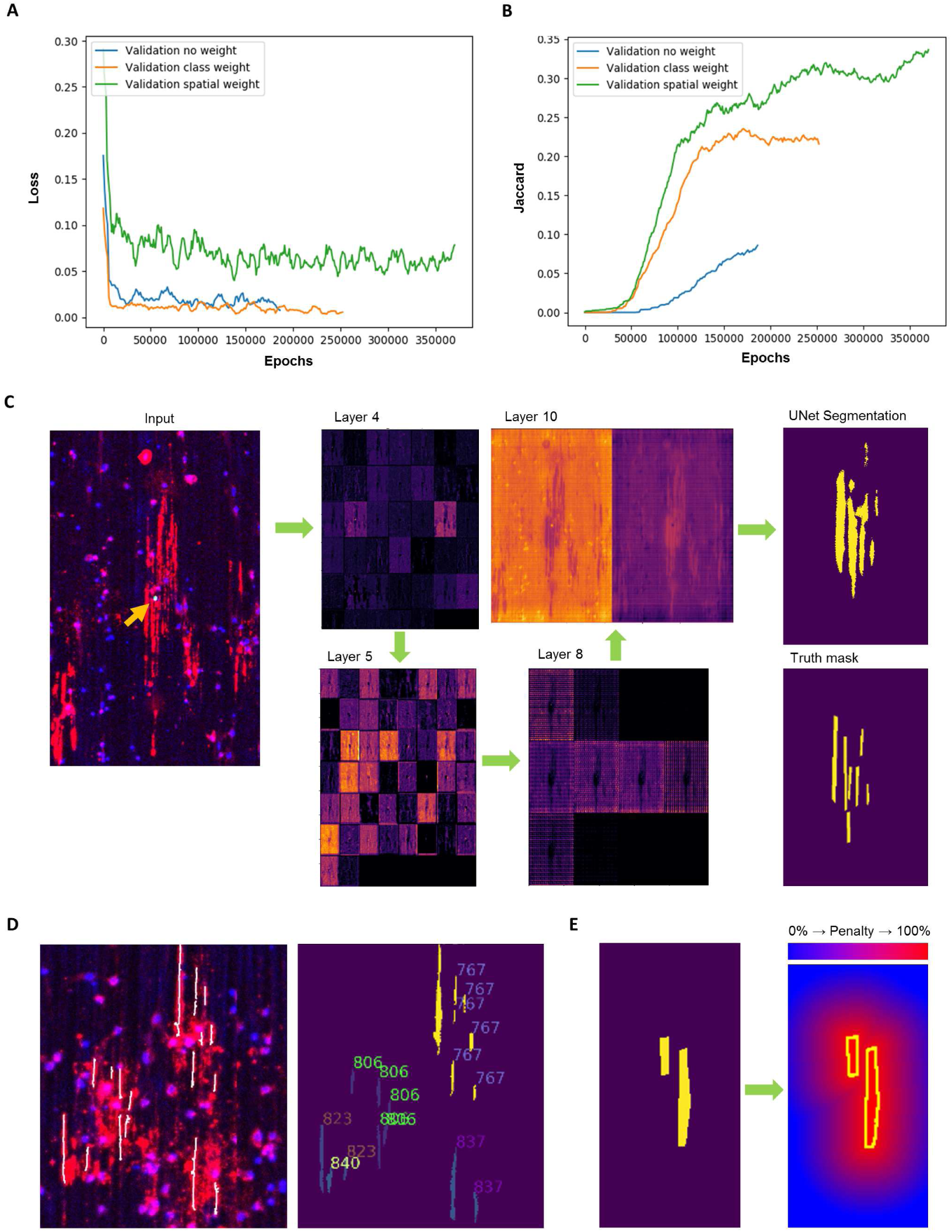
CNN training and demonstration of analysis at single-cell resolution. (A) Loss decreased continuously for all validation sets, however, the spatially weighted network did see an increase in validation loss after ∼301, 000 epochs, indicating over-fitting. Thus, the training was stopped early for testing. (B) Both class and spatial weighting improved learning rates dramatically, as indicated by an earlier and steeper increase in the JI of the validation sets. However, class weighting appears to plateau without an improvement in the JI after ∼180, 000 epochs. The spatially weighted CNN was therefore selected as our best model. (C) Sample outputs of specific layers leading to the final output “UNet segmentation”. “Layer 4” exhibited a binary response to the distribution of MBP immunoreactivity, while “Layer 5” appeared to activate for different parts of the MBP image, suggesting the extraction of features corresponding to linear ensheathments. Arrowhead shows nucleus of masked cell of interest. (D, left) Zoomed inset of the 10x whole-well image used for testing. The ensheathed fibers segmented by the UNet are overlaid (white). Hoechst is shown in blue and MBP in red. (D, right) Ensheathed fibers segmented by the UNet are annotated to indicate the specific cell they belong to, showing that the network learned to associate fibers with specific nuclei, rather than identifying individual segments globally across a whole-image. (E) Illustration of applying spatial-weighting penalization to draw “attention” to object edges during training.

To improve learning speed and accuracy, we also trained two CNNs with differently weighted loss functions. We implemented “class weighting” for the first network, whereby the loss at every pixel corresponding to ensheathments was multiplied by a factor of 10. This placed more emphasis on decisions corresponding to OL sheaths, thereby helping to “balance” the training dataset by increasing the importance of the small number of images that contained ensheathed cells. For the second network, in addition to “class weighting”, we also implemented “spatial weighting” to enhance the sensitivity of the network to learn about the edges of objects, weighing the loss function with the following exponential decay formula:

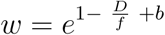

W refers to the weighted loss at each pixel, f modifies the rate of falloff from the edge, and b prevents the weights from reaching zero. D is the distance from the edge of the OL ensheathments, calculated by computing the Chebyshev distance transform of the inverted ground-truth mask. Overall, this formula weights the training loss such that it decays exponentially with distance from the margins of the ensheathed segments of interest, thus helping the network “focus” on correctly identifying the edges of objects (Figure 3E). Although both weighting schemes dramatically improved CNN performance, indicated by an earlier and steeper increase in JI, the spatially weighted network was superior, as its JI continued to improve while the class weighted network plateaued at ∼180, 000 training epochs (Figure 3B). Using these parameters, along with testing at progressive checkpoints, we found the best model was generated at ∼301, 000epochs by the spatially weighted network (Figure 3A-B). Finally, after training was performed for 350,000 epochs (about 6 days) on an NVIDIA Tesla P100-PCIE, one entire whole-well image (containing 1,667 input cells traced by a human researcher) was used to test the network for its performance on global parameters and generalizability to new data.

### 2.10 Data Availability and Statistics

The datasets generated and analysed in the current study are available from the corresponding author on request. All computational resources in this paper will be made available through deepdiscovery.org. Data are presented as mean ± standard error of the mean (SEM). Significance was set at *p<0.05, **p<0.01, ***p<0.001. Statistical analyses were completed with Graphpad Prism 5 and all Student’s t-tests are two-tailed. The log of ensheathment length was used as described [7] previously because sheath length data was log normal (log lengths were Gaussian distributions). Power analysis was performed using Lenth’s power calculator software [16, 17].

## 3 Results

### 3.1 Comparison of heuristic and deep learning algorithm segmentations

To quantify OL ensheathments, we first developed a heuristic algorithm that targeted two primary morphological properties: linearity of ensheathments, and their continuity with cell bodies (Figure 1B-C). In addition to performing qualitatively valid ensheathment segmentations (Figure 1C), this algorithm revealed the challenge of associating ensheathed segments with specific OLs, as morphological approximations of continuity and cell body size were not always viable in images containing crowded cells, faded fluorescence, or certain atypical morphologies. To resolve this, we developed a deep learning approach using a single-masked nucleus training method (See “Materials and Methods: Training”) which exhibited an enhanced capacity to associate multiple-morphological features with specific OLs (Figure 3). A qualitative visual evaluation of the output of select layers of the trained CNN confirmed that the network had learned to identify ensheathments belonging to individual OLs when prompted with images of cells with masked nuclei (Figure 3C-D). The capacity for the CNN to combine spatial and object-specific information to reliably associate specific sheaths with individual cells highlights a key advantage of our masked-nucleus training method over approximating morphological features with heuristic algorithms.

### 3.2 UNet best matched expert human accuracy

After generating heuristic and deep-learning algorithms, the performance of both systems was evaluated in relation to human researchers. Three humans were employed for this analysis and were ranked based on their experience with OL cultures—human 1 (H1) being a cell biologist, H2 a computer scientist with some experience in cell biology, and H3 a summer medical student with no previous experience working with OLs. Initial analysis of one test image containing ∼1, 000 candidate cells revealed that the output of either the heuristic or CNN algorithms fell within the range of human quantification for multiple parameters (Figure 4). No statistically significant differences were detected between the log and absolute mean sheath lengths, nor in the mean sheath length per cell (mSLC), between H1 - 3 and machine algorithms (Figure 4A–B, D). However, despite predefined segmentation criteria (see “Materials and Methods: Manual Segmentation Criteria”), variability between humans was much higher at a single-cell level (Figure 4C). Only the CNN identified a mean number of sheaths per cell that did not differ significantly from H1 (Figure 4C). The overall number of ensheathed cells identified was also consistent between groups, except for H2 (Figure 4E).

**Figure 4:**
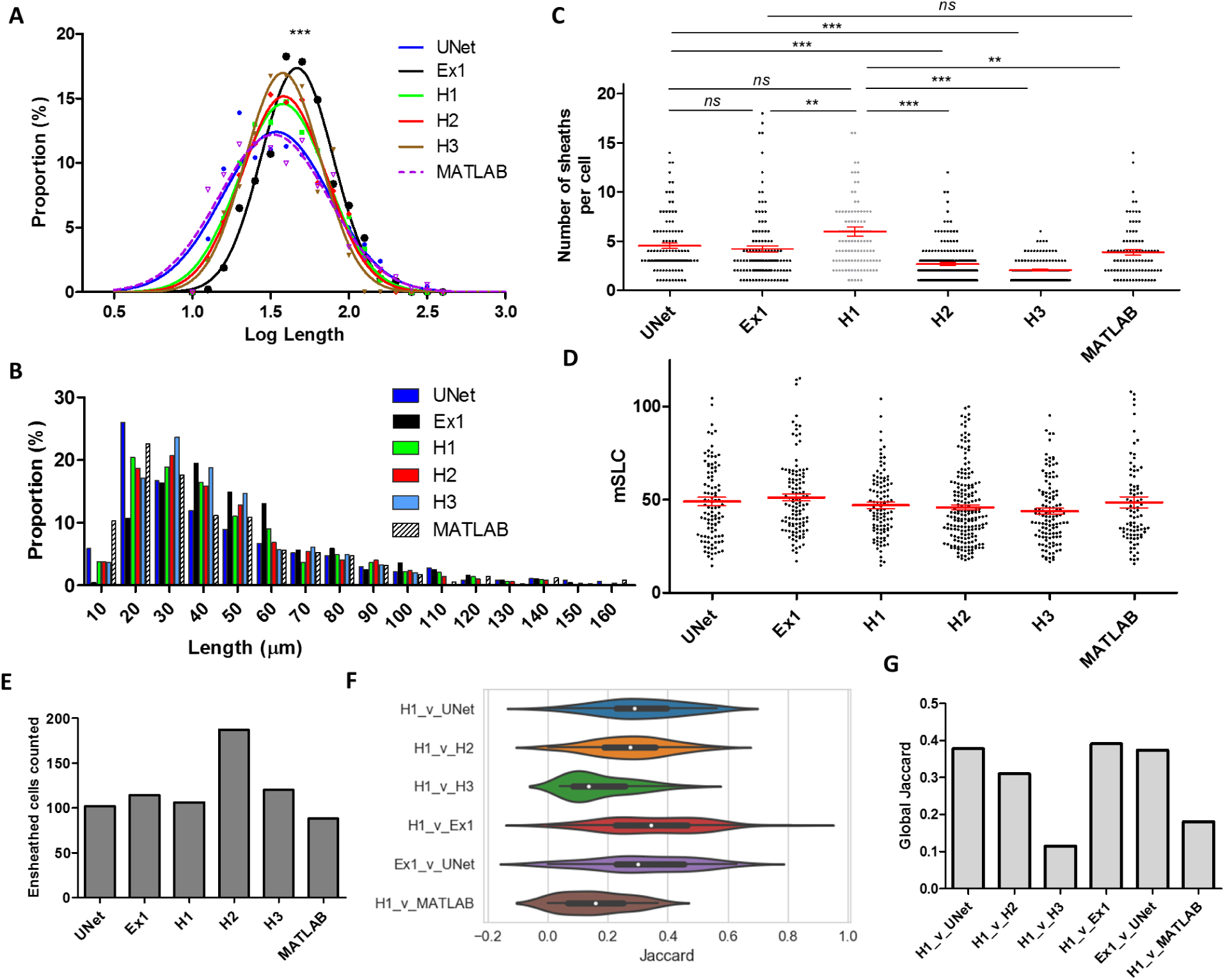
Comparison of the performance of humans (H1, H2, H3, Ex1) and machines (UNet, MATLAB) on a whole-well image. (A) Log and (B) absolute distributions of ensheathment lengths are not significantly different between machines and humans (H1 – 3) except Ex1 (one-way-ANOVA with Tukey’s post-test). (C) Only the number of sheaths per cell identified by the UNet matched that of H1 (the expert). Other humans with less expertise, and the heuristic MATLAB program, differed significantly from H1 (Kruskal-Wallis test with Dunn’s post-test). UNet also matched human Ex1, showing no bias towards training ground truth data. Error bars and midline show mean ± SEM. No significant difference between mean sheath lengths per cell (mSLC). Error bars and midline show mean ± SEM. The total number of ensheathed cells was identified similarly by all groups, except H2. This variability in analysis by human non-experts supports the need for standardized analytic systems. (F) Violin plots of the JI similarity measure when comparing the segmentations from Human 1 (H1) with H2, H3, Ex1, UNet, and the heuristic algorithm on a single-cell level. Overall, the experts best matched one another (Ex1 and H1), but the UNet also closely matched both Ex1 and H1 on a cell-to-cell basis. Middle dot of violin plots represents the median value of the distribution, and the upper/lower box limits show the upper/lower quartiles. (G) The global JI was also computed across an entire whole-well image between the same comparison groups. The UNet and Ex1 again had the highest match with H1. See also Table S1

After evaluating global morphological parameters, we then used JI comparisons to ensure that the binary algorithmic segmentations were similar to those of human researchers on a single cell and whole-image level. To assess the similarity of segmentations on individual cells, we selected the segmentations provided by H1 as the ground-truth and computed the JI for every cell that was commonly identified after pairing H1 with each researcher or computer program. We found that the CNN outperformed human participants and the heuristic algorithm in matching H1 at single-cell resolution (Figure 4F). We also computed the global JI for the entire whole-well image and found that the CNN best matched expert H1 (Figure 4G). Thus, the CNN outperforms the heuristic algorithm in analysis at single-cell resolution and in global parameters due to its heightened capacity to extract information from individual OLs.

### 3.3 UNet reduces inter-human variability and generalizes to external expert segmentations

To verify that the performance of the algorithms was not biased to the specific ground-truth used for training, we introduced an external expert (Ex1), from a different laboratory than H1, who was not involved in the development of the analytic models. Segmenting the same whole-well image, Ex1 matched the UNet on several parameters, including: the number of sheaths per cell, mSLC, single-cell JI values, global JI values, and the number of total ensheathed cells (Figure 4C–G). Ex1, however, reported a slight but significant shift of ∼4 *µ*m in mean ensheathment lengths (Figure 4A-B) and differed from H1 in identifying the mean number of sheaths per cell (Figure 4C). This difference shows that, while human experts matched each other better than the non-expert (H3), there was still slight variability in specific parameters of the analyses carried out by the two different experts. Conversely, the UNet matched Ex1 on several parameters, and even identified a mean number of sheaths per cell that was not significantly different from either expert H1 or Ex1 (Figure 4C). This capacity to match Ex1 and H1 demonstrates that the CNN is generalizable to external experts beyond the training ground-truth and highlights the capacity for automated techniques to offer an unvarying analytic standard that eliminates inherent human variability in quantification.

### 3.4 UNet performance generalizes to aligned OLs rotated at different orientations

We also assessed the capacity of the deep learning approach to quantify ensheathments at differing orientations by training a CNN with images rotated at random angles. Notably, training a CNN with rotated images dramatically decreased the training rate and resulted in a lower plateau in JI after 16 days (1,000,000 epochs) (Figure 5A). The performance of the CNN trained with rotated images was then tested with a whole well image (“UNet – rotated”), or by randomly rotating each “input” image that was cropped around a candidate cell within a whole well image (“UNet – rand rotated”). Interestingly, the random rotations of cropped “input” images during testing did not drastically affect the performance of the CNN (Figure 5). Regardless of the testing method, however, the mean length of ensheathed segments identified by the CNN trained with rotated images was significantly lower than that identified by H1 and the non-rotated control UNet (Figure 5B-C). The number of sheaths per cell, however, matched between all CNN’s and H1 (Figure 5D). As well, the CNN trained with rotated images had a slightly lower sensitivity towards ensheathed cells overall (Figure 5E), and matched H1 on global segmentations to a moderate degree (Figure 5F). Thus, our results demonstrated that a masked-nucleus training method can allow our UNet to generalize to both rotated and aligned sheaths without requiring a researcher to model more heuristic features through computer code. Notably, while training the CNN with rotated images reduced sensitivity, the network still provides a useful quantification that offers standardization to eliminate human variability in analysis.

**Figure 5:**
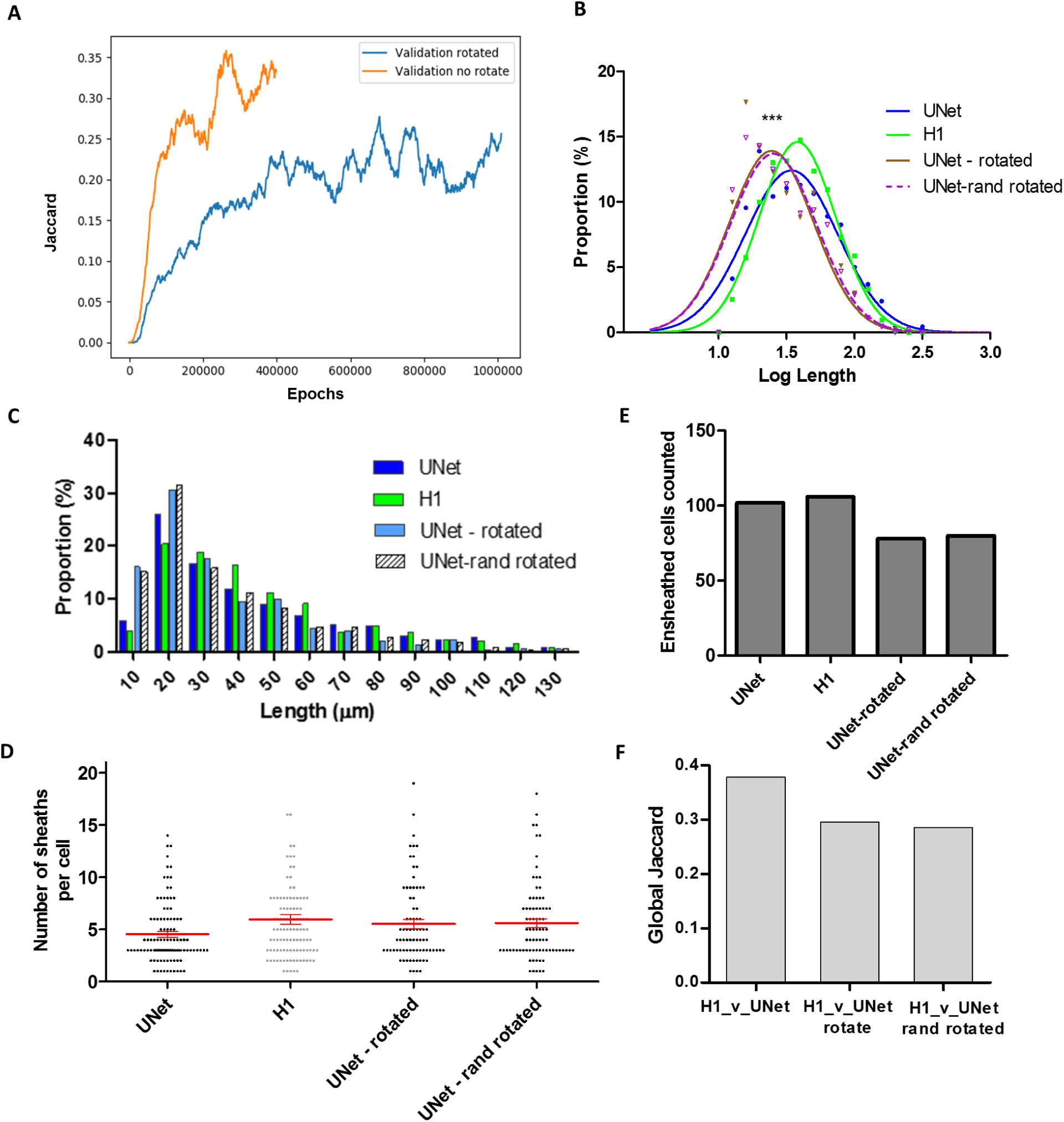
UNet performance when trained with randomly rotated images. (A) Performance of CNNs during training as monitored through JI. Compared to the control CNN (“Validation no rotate”), the JI of the CNN trained with rotated training images (“Validation rotated”) increased at a much slower rate and seemed to plateau after 16 days of training (1,000,000 epochs) at a lower JI. The UNet was then tested using a whole-well image (“UNet – rotate”) or after rotating each candidate “input” cell within a whole-well image by a random angle (“UNet – rand rotate”). The random rotation had no significant impact on testing performance. (B-C) The length distribution identified by the rotated network differed significantly from H1 and non-rotated control (UNet), but (D) the number of sheaths per cell matched between all CNN’s and H1. Error bars and midline show mean ± SEM. Additionally, (E) the CNN’s trained with rotated images also had a slightly lower sensitivity towards ensheathed cells overall, and (F) matched H1 global segmentations to a moderate degree.

### 3.5 Algorithms detect subtle difference between laminin and PDL nanofiber coatings

Having validated the analytic algorithms on both whole-image and cell-to-cell levels, we assessed the capacity of the algorithms to detect a subtle biological effect by examining the difference between PDL and laminin 1 coated nanofibers. Compared to PDL coatings, laminin has been shown to increase the number of sheaths formed per cell without significantly influencing the distribution of ensheathment lengths [7]. Both machine programs recapitulated the finding that a coating of laminin 1 increases the number of sheaths per cell compared to PDL, while identifying a novel difference in the distribution of sheath lengths, due to the increased sample size of our automated approach. For the number of sheaths formed per cell, the heuristic algorithm found a difference of 0.74 ± 0.10 sheaths (p < 0.0001) and the CNN found a difference of 0.30 ± 0.08 sheaths (p = 0.0018) between laminin and PDL coated conditions (Figure 6C). For log length distributions, both the heuristic algorithm and the CNN identified a small but statistically significant difference in means of 0.018 ± 0.0032 (p < 0.0001) and 0.0137 ± 0.0037 (p = 0.0002) log units respectively (Figure 6A-B).

**Figure 6:**
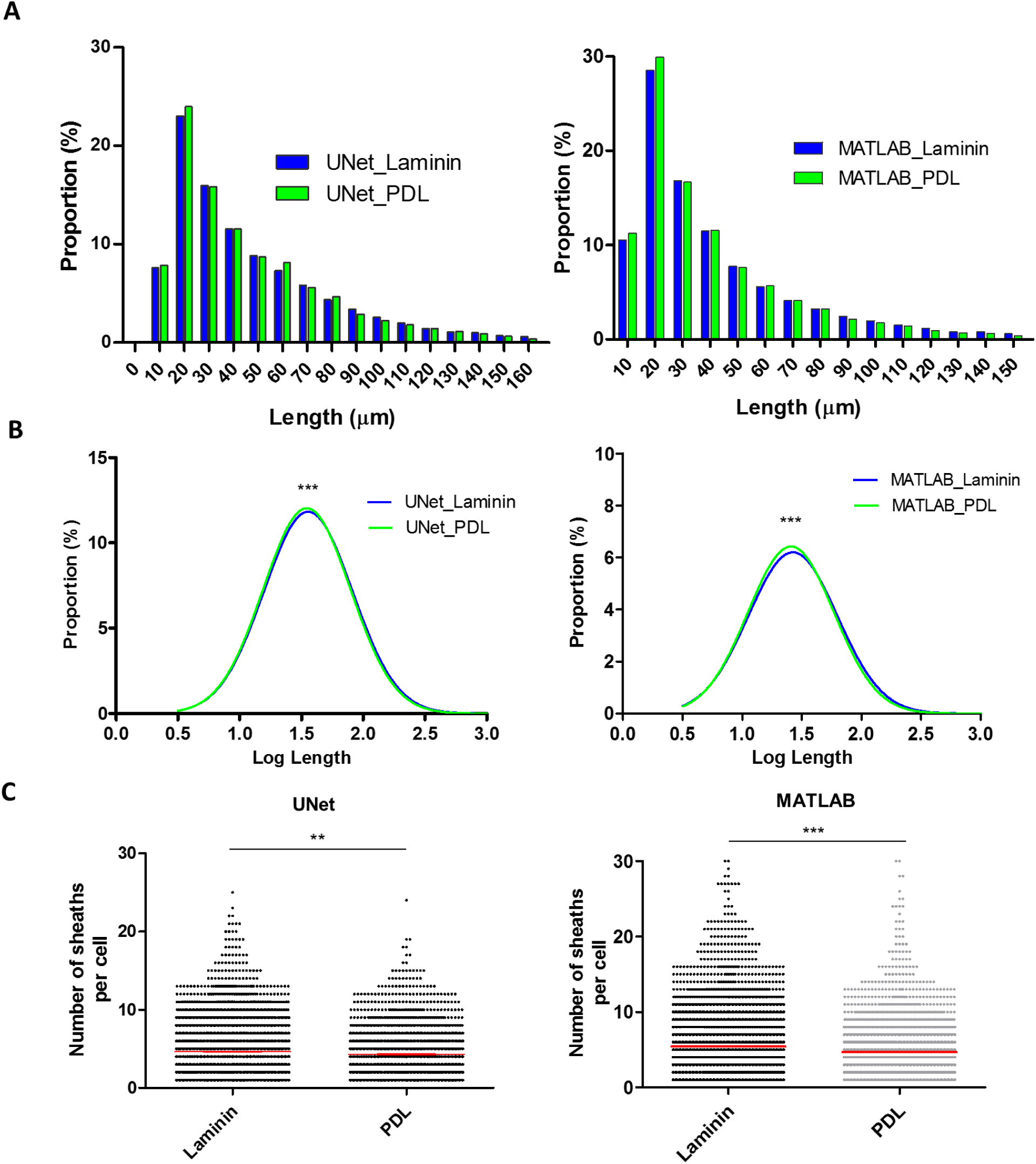
Detecting biological differences using the heuristic algorithm and CNN. (A) Absolute and (B) log length distributions had a small but significant difference in means of 0.0137 ± 0.0037 log units (p = 0.0002, Student’s T-test) as detected by the UNet, and a difference in means of 0.018 ± 0.0032 log units (p < 0.0001) as detected by the heuristic algorithmic approach. (C) Both systems detected increases in the number of sheaths formed per cell between laminin and PDL coated nanofibers. The UNet found a significant difference in means of 0.30 ± 0.08 sheaths (p < 0.0018, Mann-Whitney U-test), while the heuristic algorithm identified a significant difference in means of 0.74 ± 0.10 sheaths (p < 0.0001) between the two conditions. Cells were pooled from 6 rats and cultured as 10 duplicate wells per coating condition. Error bars and midline show mean ± SEM. See also Table S2

While Bechler et al. [7] previously observed an upwards trend in the lower and upper bounds of the log length distribution in laminin coated conditions, the difference was not statistically significant. Our results demonstrate a key advantage of the high-throughput system: by sampling ∼43 times more ensheathments in each experimental condition (∼13, 000 instead of ∼300 ensheathments), we detected a nuanced, yet significant, difference in the length of the myelin-like ensheathments in the laminin 1 coated condition. A power analysis, using Lenth’s Power Calculator [16, 17], demonstrates that a sample size of 13,000 sheaths (from 10 wells with ∼260 ensheathed cells and ∼5 sheaths per cell) can detect a difference of 0.01 log lengths with 80

### 3.6 Automated pipeline permits high-throughput assessment of OL ensheathment

Finally, we evaluated the speed of our analytic algorithms for practical use. Our external expert (Ex1) took ∼2 hours to trace all MBP stained ensheathments (corresponding to ∼100 ensheathed cells) in a single well. Assuming that a researcher traces for about 8 hours per day, it would take approximately 24 days to analyze an entire plate. In contrast, the heuristic algorithm and CNN (using a NVIDIA GTX 1070 graphics card) analyzed the entire surface of a 96-well plate in ∼1 day (Table 1). Automated microscopy also reduced the imaging time from approximately 77 manual hours to 15 automated hours for a full plate (Table 2), with imaging performed autonomously overnight after setting up and calibrating the system. Altogether, the automated pipeline dramatically enhanced the speed of analysis while largely eliminating manual-labor, thus providing the capacity to screen hundreds of compounds within a week.

**Table 1:** comparison of human and machine approaches for quantifying OL ensheathment

|  | Time per cell<br>(seconds) | Time per well<br>(mins) | Time 96 wells<br>(days)**** | Benefits | Limitations |
| --- | --- | --- | --- | --- | --- |
| Human | 80 | *400 | ~26.7 | - Adaptable<br>- Fast learner | - Variable<br>- slow and costly |
| MATLAB | - | **20 | ~1.3 | - Fast<br>- Standardization<br>- User customizability | - Approximates<br>cell-by-cell analysis |
| UNet | 1.5 | ***50 | ~3.3 | - Moderate-Fast speed<br>- Adaptable given time<br>to train<br>- Standardization<br>- Analysis at level of<br>single-cells | - Training time long<br>- Uncertain<br>generalizability |
| UNet (GPU<br>accelerated) | 0.5 | ***15 | ~1 |  |  |

\*assume ~300 ensheathed cells per well
\*\*program cuts whole-wells into 16 sections if insufficient memory, taking ~1 minute on each section
\*\*\*assume ~2000 candidates per well
\*\*\*\*analysis time similar for 384-well plates due to identical plate area

**Table 2:**
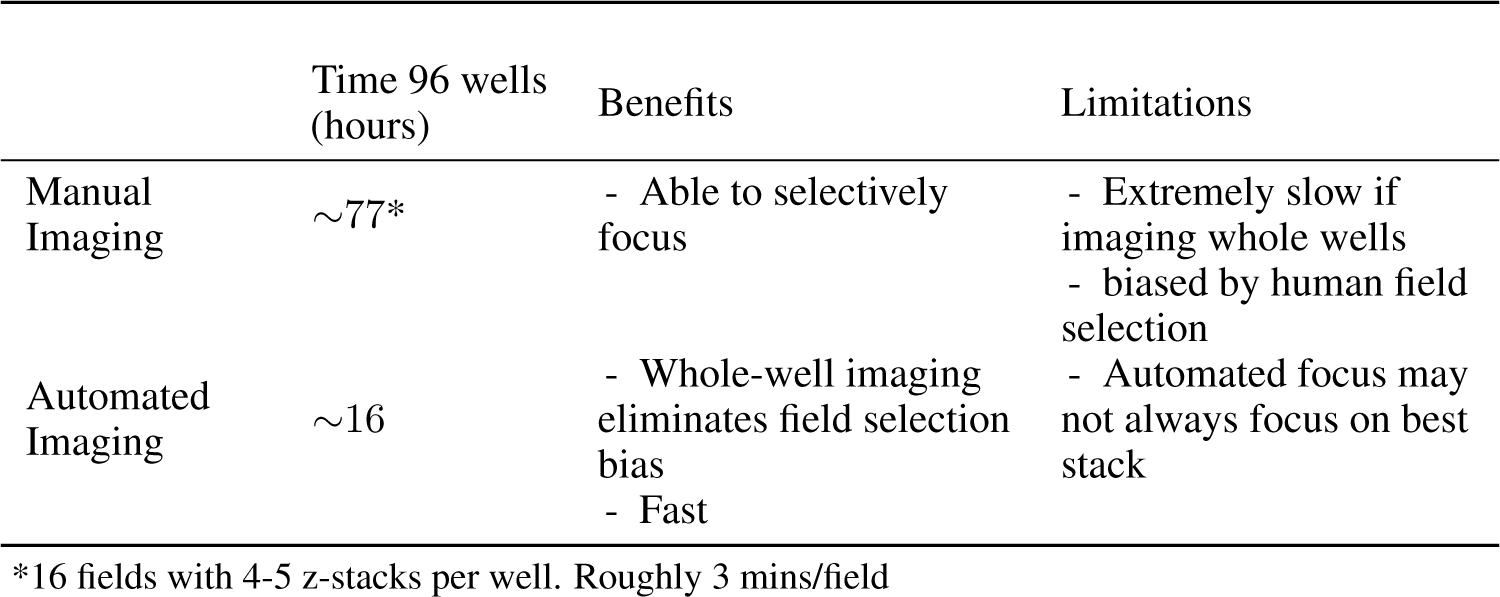
comparison of imaging time. Automated imaging approach reduces manual labor significantly, while also reducing capture time due to automated focus and pre-defined acquisition area

## 4. Discussion

To address the need for high-throughput systems that can quantify multiple morphological properties of OLs and associate them with specific cells, we developed an analytic pipeline using nanofiber cultures, automated imaging, and quantitative algorithms. We demonstrate that this analytic paradigm has the capacity to detect finely nuanced biological differences and to match the quality of quantification carried out by human experts.

The biological and computational techniques employed in our experimental system have unique advantages and limitations. The reduced complexity of the nanofiber culture system, which lacks axons and other factors that may influence sheath stability and extension [18, 19, 20], is advantageous to isolate OL intrinsic responses while also substantially increasing the reproducibility of quantification for high-throughput screening. OLs cultured on nanofibers broadly recapitulate the temporal and spatial progression of differentiation seen in vivo [6], forming compact myelin-like MBP positive layered sheaths that wrap nanofibers [7]. Further, the parallel arrangement of the nanofibers facilitated the development of our analytic programs, as it was not necessary to disentangle myelin-like sheaths crossing-over one another, as occurs in typical axon-OL co-cultures. Nanofiber cultures are also highly scalable, allowing many different conditions to be analyzed in parallel.

A final advantage of the nanofiber system is that it facilitates the assessment of multiple biologically relevant parameters, such as the number of sheaths formed per cell, the distribution of sheath lengths and widths, and the proportion of cells that form ensheathments [2, 21]. Access to quantitative information for multiple parameters per cell provides key insight into which cellular process may be affected by any given treatment. For example, changes in the number of ensheathed cells implies an effect on OPC proliferation/differentiation, whereas changes in sheath lengths would suggest altered process extension or stabilization. Acquiring multiple parameters may reveal unexplored correlations, such as between the mean number of sheaths per cell and sheath length per cell. In contrast, other existing high-throughput systems often trade multi-parametric analysis for speed. For example, a multi-well assay that measures the MBP intensity of OL ensheathments around single micropillars obtains a high-throughput level of analysis, but discards information pertaining to sheath length and number [8]. The automated pipeline we developed retains analytic speed while extracting multiple parameters from individual OLs, allowing the detection of nuanced physiological processes that can then be further studied in more complex systems.

Both computational approaches presented offer advantages beyond reducing human workloads and extracting multiple morphological features. For example, both algorithms provide standardized reproducible analysis, eliminating the human variability that may result from internal biases, mistakes, and even environmental factors, such as screen brightness, background lighting, and time-constraints. Analytic speed may be enhanced nearly indefinitely by employing parallel computing workstations and faster GPUs. Limiting factors become the time for sample preparation, rather than daunting hours of manual analysis. Finally, the architectural design of the CNN also offers a significant advantage. Compared to the original UNet architecture proposed by Ronneberger et al., which scaled to 1024 filters in the deepest layers [10], our reduced network structure is computationally more efficient, scaling to only 50 filters in the deepest layer, allowing our network to run in reasonable time on common CPUs.

Aside from similar advantages, the two programs differed in their accuracy and speed. The heuristic approach is limited as it uses computational approximations to extract global parameters from an image and then attempts to mimic cell-by-cell analysis by associating ensheathments with nearby nuclei. In contrast, the CNN provides a more “human-like” assessment of nanofibers by performing high-level decisions using spatial and object-specific information simultaneously to associate specific parameters with individual cells. Although the heuristic algorithm is less accurate in this capacity, it is important to highlight that the approach is still consistent, fast, and responds more readily to user-specified parameters. The heuristic algorithm is particularly appropriate for studies composed of insufficient data to train a neural network, or for analyzing data-sets containing high-variability that require rapid optimization of analytic algorithms.

A final consideration is the generalizability of these programs. While the heuristic algorithm was designed explicitly for aligned nanofiber cultures, it has the immediate capacity to analyze other linear sheath-like objects. The major obstacle to this generalizability is that such an application may require a researcher to have programming experience to alter the program to fully suit their needs. In contrast, the CNN can match human segmentations on a variety of computational problems without substantially altering the algorithm. For example, we demonstrated that the masked-nucleus training method can generate a UNet capable of segmenting aligned OLs in multiple orientations (Figure 5), suggesting that this technique may be applied to more complex OL-neuron co-culture systems. Given that the CNN performs well with evenly-distributed parallel sheaths in our nanofiber culture system, this quantitative approach could be reasonably applied to tissue sections from CNS regions with an aligned geometry. The major limiting factor for generalizability with the CNN is the time required to re-train the network and to create ground-truth data for supervised learning. While training is ideally only performed once, and we trained the network with a variety of cell culture and imaging parameters, the full capacity of the CNN to generalize across larger variations in experimental preparations and imaging systems remains to be determined.

## 5 Conclusion

In this study, we have developed a powerful paradigm for high-throughput screening of OL ensheathments through the combination of state-of-the-art deep learning technology and readily available imaging and cell culture systems. Our methods extract multiple morphological parameters from individual OLs, preserve human analytic quality, remove bias and variability, and facilitate massive increases in sample size and analytic speed. These new developments promise to advance the understanding of fundamental OL physiology and accelerate the discovery of new drugs for the treatment of demyelinating diseases.

## Supporting information

## 6 Acknowledgments

We thank Compute Canada for access to computer clusters to train the neural network, and Drs. Thomas Stroh and Liliana Pedraza at the Microscopy Core at the Montreal Neurological Institute for technical assistance. This research was undertaken thanks in part to funding from the Canada First Research Excellence Fund, awarded to McGill University for the Healthy Brains for Healthy Lives initiative. Additional support was provided by grants from the International Progressive Multiple Sclerosis Alliance (BRAVEinMS) and the Multiple Sclerosis Society of Canada.

