## Supplementary material for "Deep Learning for High-Throughput Quantification of Oligodendrocyte Ensheathment at Single-Cell Resolution"

### SUPPLEMENTARY METHODS AND DATA

**Table S1: Average sheath lengths with standard deviation.**

Since the sheath length data was log normal (log lengths followed a Gaussian distribution), the average log length and standard deviation were used for statistical tests. The upper and lower quartile of the distribution are also reported in microns, along with the number of sheaths identified per condition.

| Related figure | Condition | Mean log(length) | Standard deviation log(length) | Mean length (microns) | Number of Sheaths | 25th Percentile (microns) | 75th Percentile (microns) |
|---|---|---|---|---|---|---|---|
| 3 A, B | UNet | 1.58 | 0.29 | 48.84 | 461 | 21.63 | 63.94 |
| 3 A, B | H1 | 1.60 | 0.25 | 47.25 | 631 | 25.34 | 59.89 |
| 3 A, B | H2 | 1.60 | 0.25 | 47.58 | 497 | 26.69 | 57.68 |
| 3 A, B | H3 | 1.58 | 0.22 | 42.47 | 245 | 26.68 | 53.17 |
| 3 A, B | MATLAB | 1.57 | 0.30 | 47.85 | 341 | 20.93 | 59.81 |
| 3 A, B | Ex1 | 1.68 | 0.23 | 54.25 | 477 | 33.75 | 65.83 |

**Table S2: Average sheath lengths with standard deviation.**

Since the sheath length data was log normal (log lengths followed a Gaussian distribution), the average log length and standard deviation were used for statistical tests. The upper and lower quartile of the distribution are also reported in microns, along with the number of sheaths identified per condition.

| Related figure | Condition | Mean log(length) | Standard deviation log(length) | Mean length (microns) | Number of Sheaths | 25th Percentile (microns) | 75th Percentile (microns) |
|---|---|---|---|---|---|---|---|
| 5 A, B | UNet, Laminin | 1.597 | 0.31 | 51.43 | 18512 | 22.27 | 66.17 |
| 5 A, B | UNet, PDL | 1.583 | 0.30 | 49.1 | 10490 | 21.71 | 63.23 |
| 5 A, B | MATLAB, Laminin | 1.53 | 0.30 | 44.92 | 23264 | 19.16 | 54.87 |
| 5 A, B | MATLAB,PDL | 1.52 | 0.30 | 42.33 | 13356 | 18.76 | 52.08 |

### Supplementary cell culture and staining methods

P2 Sprague-Dawley rat brains were isolated by dissection in ice cold HBSS, the meninges removed, and cortices isolated. These were minced and digested at 37˚C in 0.025% trypsin-EDTA and 0.2 mg/mL DNAseI for 20 and 3 min respectively. Dissociated cells were plated in PDL-coated T75 flasks (5 µg/mL) in DMEM containing 10% FBS and 1% pen/strep, and cultured for 2-3 weeks at 37˚C with 5% $CO_2$ and full media changes every 2 days. In these conditions, glial populations rapidly expand, permitting isolation from other cell types based on differential adhesion. Media in the flask was replaced with 37˚C 0.01% trypsin-EDTA in HBSS for 5 min, then swirled and aspirated to remove weakly-adherent microglia. DMEM was then added and the flasks were vigorously hit 20 times with a styrofoam box. This mechanical agitation detached oligodendrocyte precursor cells (OPCs) while leaving most astrocytes attached. The media-cell suspension was plated in uncoated Petri dishes and incubated for 30 min to further purify the cells[6, 22], as astrocytes and microglia adhere more readily than OPCs. The OPC-containing supernatant was then collected and centrifuged for 5 min at 677 x g using a swinging bucket rotor in a Centra CL2 centrifuge (Thermo). The cell pellet was resuspended in Sato media (below), and triturated through 18 and then 22 gauge needles. The suspension was strained through a 70 µm filter, stained with Trypan blue to assess viability, and counted using a haemocytometer. After isolation, cells were cultured in modified OL-defined differentiation media (DMEM, 5 µg/ml insulin, 100 µg/ml transferrin, 30 nM sodium selenite, 30 nM triiodothyronine, 6.3 ng/ml progesterone, 16 µg/ml putrescine, 100 U/ml penicillin, 100 µg/ml streptomycin, and 2 mM glutamax). After the stated culture period, live cells were then incubated with the O4 antibody for 15 min at 37$^{o}$C and then fixed by adding 100 µL of chilled 4% paraformaldehyde in 20% sucrose to each well for 15 min at RT. For MBP staining, fixed cells were blocked and permeabilized for 15 min with 5% HINHS (heat-inactivated normal horse serum) in 0.1% Triton X-100 at RT, and then incubated with a primary chicken anti-MBP antibody in 5% HINHS (1:1000; Aves labs) for 4 hr at RT. The OLs were then washed in PBS before being incubated with IgM Cy3 for O4 stained cells, or Alexa-546 conjugated goat anti-chicken secondary antibody (1:1000; Molecular Probes) for anti-MBP stained cells, in addition to 1 µg/mL of DNA staining Hoechst 33342 (Sigma) for 2 hr at RT in 5% HINHS. Cells were washed again with PBS and kept at 4˚C until they were imaged.
